# SadB acts as a master regulator modulating *Pseudomonas aeruginosa* pathogenicity

**DOI:** 10.1101/2025.09.11.675511

**Authors:** Maria Papangeli, Jeni Luckett, Stephan Heeb, Morgan R. Alexander, Paul Williams, Jean-Frédéric Dubern

**Author notes:** Department of Infectious Disease, Imperial College, London, UK.

## Abstract

*Pseudomonas aeruginosa sadB* mutants exhibit a hyper-swarming, biofilm defective phenotype. To acquire insights into the *in vivo* contribution of *sadB* to virulence in a mouse soft tissue infection model, we employed bioluminescence imaging and histopathology. Compared with the parent PAO1 strain, the Δ*sadB* mutant was highly attenuated and rapidly cleared from the infection site whereas genetic complementation conferring constitutive expression of *sadB* resulted in a much more persistent phenotype. Transcriptome analysis of exponential and stationary phase planktonic cells revealed that SadB modulates expression of diverse genes involved in biofilm development, quorum sensing (QS), secondary metabolite production, iron acquisition, virulence, protein secretion and anaerobiosis. In Δ*sadB,* we observed log phase induction of the *rhl* and *pqs* QS systems, increased production of siderophores and pyocyanin, differential regulation of genes involved in c-di-GMP signalling and a growth defect under static conditions. Since Δ*sadB* is a hyperswarmer and as swarming requires rhamnolipids, this suggested that deletion of *sadB* may impact on the timing and level of rhamnolipids produced. In Δ*sadB*, the *rhlA* and *rhlB* genes were induced in log rather than stationary phase resulting in overproduction of rhamnolipids. Since they also contribute to biofilm maturation and dispersal and can act as anti-adhesives, we deleted the *rhlA* in Δ*sadB* and observed that biofilm formation was restored offering mechanistic insight into the biofilm defective phenotype of Δ*sadB*. SadB clearly has a more global role than previously appreciated and acts as a master regulator of diverse genes involved in environmental adaptation, biofilm formation and virulence.

**IMPORTANCE:** Biofilms are characterised by their intrinsic tolerance to antibiotics, host immune defences and ability to cause persistent infections. In *Pseudomonas aeruginosa,* mutation of the surface attachment defect gene*, sadB* results in cells that are biofilm, defective, hyperswarmers. Here we describe an investigation of the contribution of SadB to virulence. Our study indicates that in *P. aeruginosa* a *sadB* deletion mutant is greatly attenuated in an acute mouse skin infection model. We further demonstrate that SadB acts as a pleiotropic regulator of many genes involved in biofilm development, quorum sensing, iron acquisition, protein secretion and anaerobiosis, uncovering a wider role in pathogenesis than formerly recognized, and that it has considerable potential as a novel protein target for antibacterial drug discovery.

## INTRODUCTION

*Pseudomonas aeruginosa* is an opportunistic human pathogen that produces an armoury of cell-associated and extracellular virulence determinants controlled via global regulatory systems operating at both transcriptional and post-transcriptional levels (1). It can adopt either planktonic free living or sessile biofilm-associated lifestyles which respectively contribute to acute and chronic infections. *P. aeruginosa* is capable of flagella-mediated swimming through liquids or attaching and migrating over surfaces via swarming or type IV pilus-mediated twitching motility (2). Both flagella and type IV pili can also act as adhesins and mechanosensors facilitating biotic and abiotic surface colonization (3). With respect to biofilm development, the first committed step involves the transition from reversible (attachment via the cell pole) to irreversible surface attachment (via the long axis of the cell) (2, 4). This in turn leads to micro- and macro- colony formation and biofilm maturation during which the bacteria become embedded within an extracellular matrix consisting of exopolysaccharides, extracellular DNA, lipids and proteins (5, 6, 7).

Biofilm formation and swarming motility in *P. aeruginosa* are inversely regulated. One of the first *sad* (<u>s</u>urface <u>a</u>ttachment <u>d</u>efect) genes discovered to play a role in this regulatory pathway was *sadB.* A transposon insertion in the *P. aeruginosa* strain PA14 *sadB* gene enhanced swarming but prevented biofilm formation (8, 9). Conversely, overexpression of *sadB* promoted biofilm development but reduced swarming motility. Further work resulted in a model for *P. aeruginosa* PA14 in which the SadB-dependent inverse regulation of swarming and biofilm was proposed to be mediated via viscosity-dependent modulation of flagellar reversal rates and to influence Pel exopolysaccharide production via a mechanism involving the *pil-chp* chemotaxis pathway (9, 10).

In diverse bacterial species the transition between motile and sessile lifestyles is controlled via pathways involving the second messenger cyclic-di-guanylate (c-di-GMP) whereby elevated c-di-GMP levels promote biofilm formation but inhibit motility (11, 2). C-di-GMP signalling depends on the production and turnover of this second messenger via diguanylate cyclases (DGCs) and phosphodiesterases (PDEs) respectively while cellular responses are mediated via specific c-di-GMP binding receptors (12). High levels of c-di-GMP increase production of key *P. aeruginosa* biofilm components including Pel, Psl and alginate exopolysaccharides via transcriptional and post-transcriptional regulatory pathways. In *P. aeruginosa* PA14, deletion of the cyclase gene, *sadC* resulted in a similar biofilm defective, hyper-swarming phenotype to that of a *sadB* mutant which could be restored to the WT phenotype by *sadB* overexpression suggesting that *sadB* acts downstream of *sadC* (13). C-di-GMP produced via SadC is degraded via the PDE, BifA, which together with SadC, inversely regulates biofilm and swarming motility in a *sadB* dependent manner in *P. aeruginosa* PA14. BifA is considered to act upstream of *sadB* (14) which is highly conserved amongst the pseudomonads but is absent from most other bacterial species*. sadB* codes for a cytoplasmic protein (9) which, in *Pseudomonas fluorescens* F113 has been reported to bind c-di-GMP (15). However, more recently, (16) reported that SadB from *P. aeruginosa* functions as an anti-adaptor protein, post-translationally regulating AmrZ, a member of the ribbon-helix-helix family of DNA binding proteins that acts as a transcriptional activator and repressor of multiple genes involved in virulence and biofilm formation (17; 18).

Here we confirm that a *P. aeruginosa* PAO1 *sadB* deletion mutant exhibits the same biofilm defective, hyper-swarming phenotype observed in strain PA14, is highly attenuated in a mouse infection model and acts as a pleiotropic regulator of diverse genes involved in biofilm development, quorum sensing, secondary metabolite production, iron acquisition, virulence, protein secretion and anaerobiosis. Biofilm development by Δ*sadB* could be restored by deleting *rhlA* consistent with the overproduction of rhamnolipids observed in the deletion mutant contributing to our understanding of the inability of *P. aeruginosa sadB* mutants to form mature biofilms.

## RESULTS

### A *P. aeruginosa* PAO1 Δ*sadB* mutant is highly attenuated in a mouse wound infection model

To determine whether mutation of *sadB* had a similar impact on swarming and biofilm formation in *P. aeruginosa* PAO1 to that described for strain PA14 (10), we deleted the PAO1 *sadB* gene (Table S1). PAO1 Δ*sadB* is a hyper-swarmer (Fig. 1A), fails to form a mature biofilm (Fig. 1B and 1C) and does not express *sadB* (Fig. 1D) but can be complemented genetically by introducing *sadB* on a plasmid expression vector (Fig. 1D). To evaluate the contribution of SadB to colonization and persistence in a subcutaneous mouse infection model, we constructed bioluminescent *P. aeruginosa* PAO1 WT and Δ*sadB* mutants by introducing a CTX::*tac’*-*luxCDABE* fusion onto the chromosome to avoid loss during infection that accompanies the use of plasmid expression vectors. In addition, we complemented the Δ*sadB* mutant with a CTX::*tac’-sadB-luxCDABE* fusion that constitutively expresses *sadB* (Table S1). The phenotype of each bioluminescent strain with respect to *sadB* expression and swarming was confirmed (Fig. S1).

**FIG 1.**
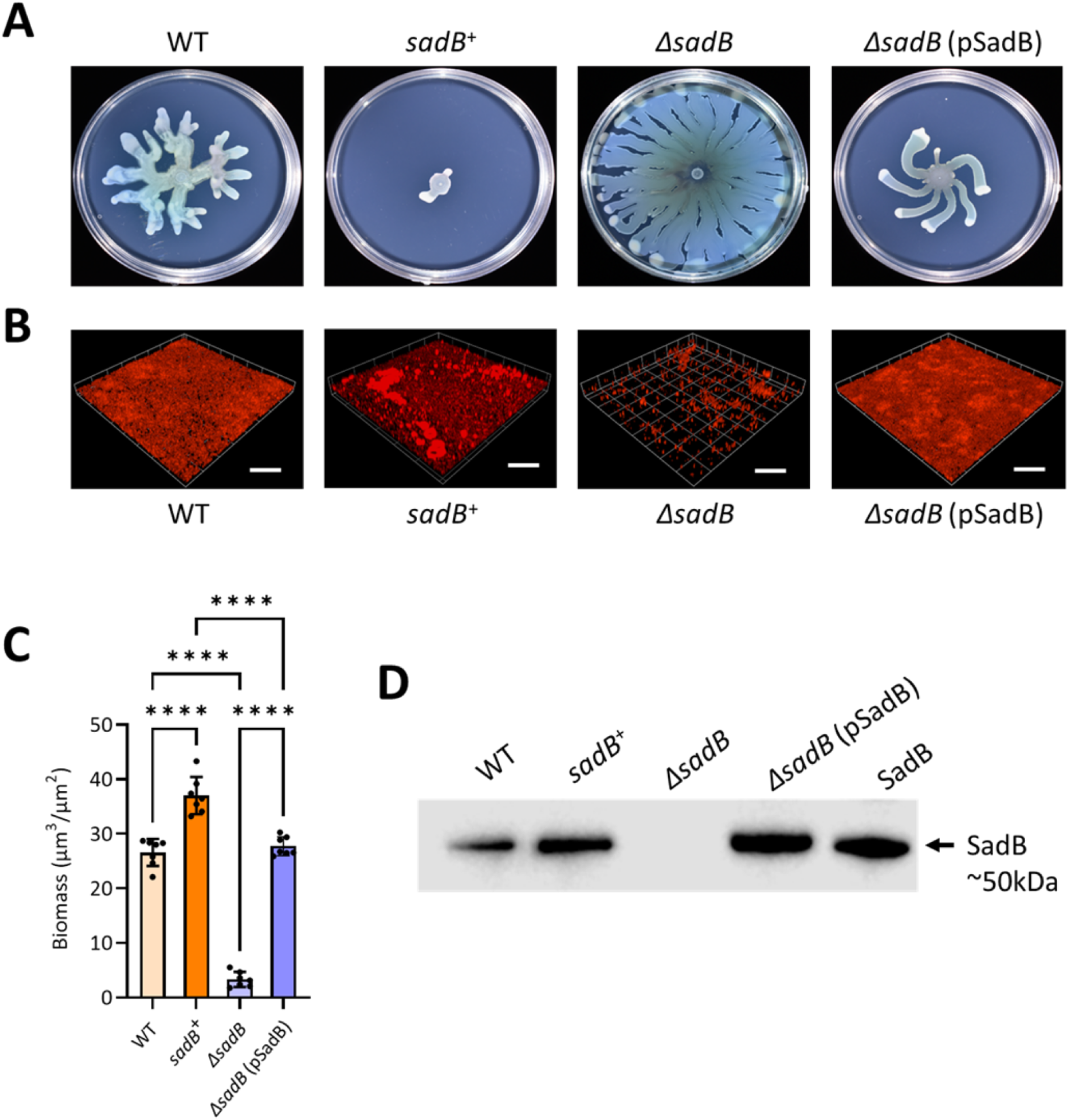
Swarming and biofilm phenotypes of *P. aeruginosa* strains lacking or constitutively overexpressing *sadB.* (A) Representative images of swarming motilities displayed by *P. aeruginosa* PAO1 WT compared with *sadB*^+^, Δ*sadB* and Δ*sadB* genetically complemented with pSadB. (B) Confocal images of biofilm formation by the same strains as in (A) after 48 h incubation in RPMI-1640 medium. Biofilms were stained with FM4-64 dye. Scale bar: 100 µm. (C) Quantification of biofilm biomass from confocal images shown in (B). Statistical differences between group means were determined by one-way ANOVA analysis using Tukey’s multiple comparisons test (\*\*\*\**p* < 0.0001). (D) Detection of SadB in cytoplasmic extracts of the *P. aeruginosa* strains as in (A) by immunoblotting compared with the purified recombinant SadB protein.

The impact of deleting *sadB* on *P. aeruginosa* growth and survival *in vivo*, was investigated using a simple and reproducible mouse model that closely mimics human cutaneous wound infections without requiring high bacterial inocula or traumatic injury (19) that could be followed in real time by live animal luminescence imaging (20). Each of the three *P. aeruginosa* strains (1×10^5^ cells) was mixed with Cytodex1 beads and injected subcutaneously. The progress of infection was tracked by imaging for metabolically active bioluminescent bacteria each day for 4 days. Fig.2 shows that the WT established within the dermis inoculation site whereas the Δ*sadB* mutant was unable to colonize. Compared with the parent strain, no bioluminescence was detectable suggesting that no metabolically active Δ*sadB* cells were present after one day post-infection (Fig. 2A-C). Furthermore, no viable bacteria could be isolated from the Δ*sadB* infection site (Fig. S2). Genetic complementation of the Δ*sadB* mutant in which *sadB* expression was driven via a constitutive p*tac* promoter restored the ability of *P. aeruginosa* to establish infection and to colonize a much larger infection site than the parent, WT strain. (Fig. 2A-D).

**FIG 2.**
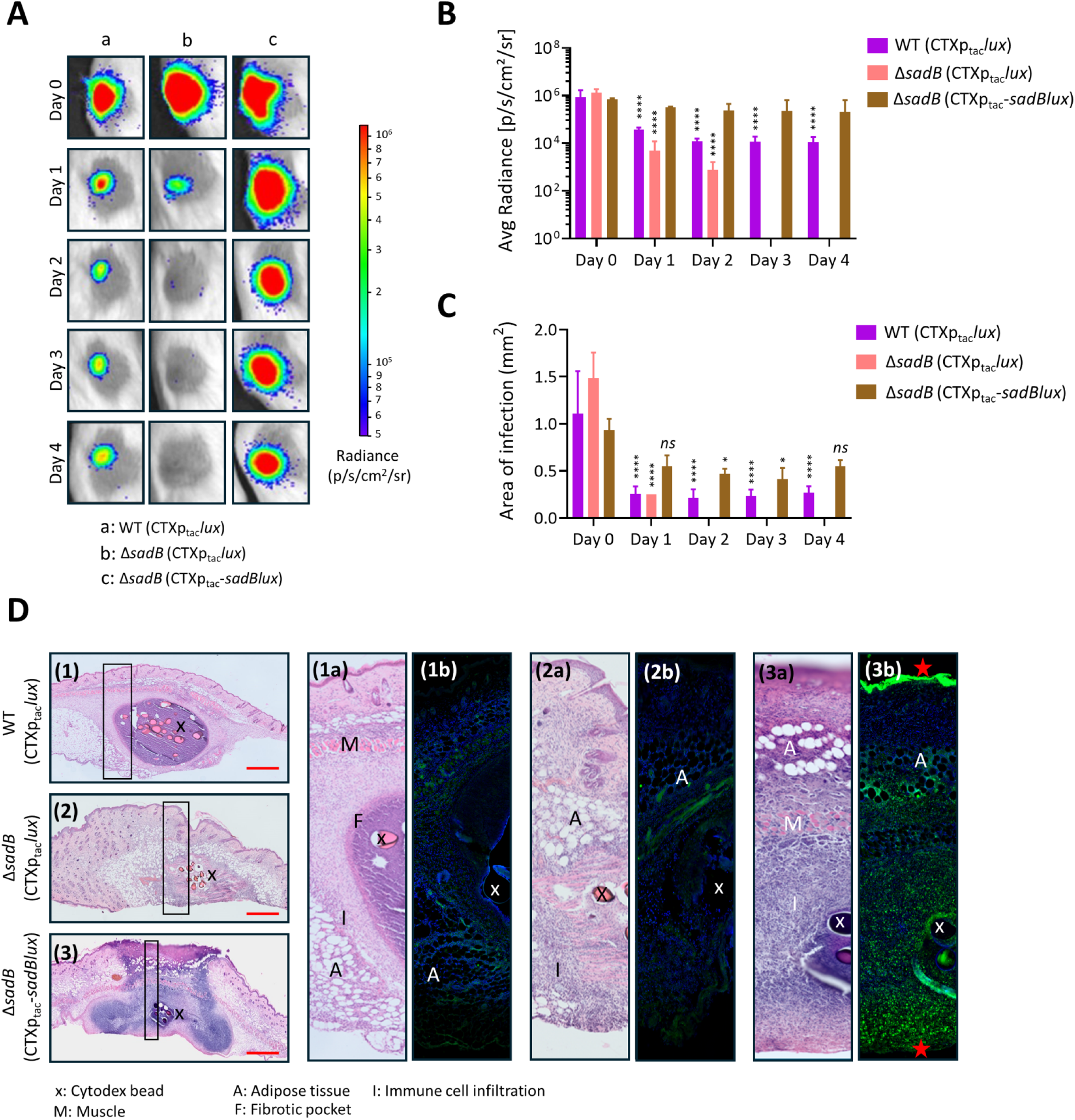
*P. aeruginosa* Δs*adB* is highly attenuated in a mouse subcutaneous infection model compared with WT and *sadB*^+^. (A) Luminescent images of the infection sites in live mice over 4 days after inoculation with bioluminescent *P. aeruginosa* strains (WT (CTX::*tac’-luxCDABE*), Δ*sadB* (CTX::*tac’-luxCDABE*) and Δ*sadB* (CTX::*tac’-sadB-luxCDABE*). n=3 mice per bacterial genotype were imaged using an IVIS Spectrum after inoculation (day 0) and on each subsequent day for 4 days. (B) Quantification of metabolically active bacteria and (C) quantification of the area of infection for each strain imaged. Statistical differences between group means were determined by Two-way ANOVA analysis using Tukey’s multiple comparisons test to day 0 (\**p* < 0.05, \*\*\*\**p* < 0.0001). (D) Histological assessment of *P. aeruginosa* infection site tissue. Panels (1), (2) and (3) show haematoxylin and eosin staining of excised tissues from mice infected respectively with *P. aeruginosa* WT, Δ*sadB* and Δ*sadB* (CTX::*tac’-sadB-luxCDABE*). The boxed regions in each panel are shown as enlarged panels in (1a), (2a) and (3a). *P. aeruginosa* WT infection established within a fibrotic pocket (panels (1) and (1a)) whereas no defined fibrotic pockets were apparent following Δ*sadB* (panels (2) and (2a)) or Δ*sadB* (CTX::*tac’-sadB-luxCDABE*) (panels (3) and (3(a)). For panels (1b), (2b) and (3b), bacterial cells were visualized by IHC using polyclonal antibodies raised against *P. aeruginosa* (green) and tissue sections counterstained for DNA with POPO-1 (blue). Panel 3(b) shows that for Δ*sadB* (CTX::*tac’-sadB-luxCDABE*), bacterial cells were present on the dermal surface and in clusters at the interface between the mouse skin and the underlying tissue (see red stars). In addition, although no fibrotic pocket formed there is evidence of leukocyte infiltration in 3(b). A, adipose tissue; M, muscle; F, fibrotic pocket; I, Immune cell infiltration; Cytodex beads (x).

At day 4, the mice were euthanized and the intact infection sites dissected, formalin fixed and embedded in wax. The architecture and spatial localisation of the infection site was assessed by comparing parallel tissue sections of each site, one stained with haematoxylin and eosin and the other used for immunohistochemistry (IHC) to localize the *P. aeruginosa* cells in each infection site. Histopathological assessment of the dermis infection site revealed marked strain dependent differences in tissue architecture (Fig. 2D). The WT underwent encapsulation within a discrete fibrotic pocket (into which leukocyte infiltration was observed). In contrast, for Δ*sadB*, no fibrotic pocket formed although some leukocyte infiltration was visible. Inoculation with the Δ*sadB* mutant constitutively expressing *sadB* did not lead to the formation of fibrotic pockets and stimulated the marked infiltration of leukocytes within and beyond the infection site into the adipose and subdermal tissue layers. IHC was used to localize *P. aeruginosa* WT, Δ*sadB* and Δ*sadB* CTX::*tac’-sadB-luxCDABE* cells within, as well as outside, the excised and sectioned infection sites. These images (Fig. 2D, (1b), (2b) and (3b) revealed the presence of WT bacteria present both inside and outside the fibrotic pocket within the infection site (Fig. 2D (1b)). Few Δ*sadB* cells were only detected within the infection site (Fig. 2D (2b)) while Δ*sadB* CTX::*tac’-sadB-luxCDABE* cells were observed within and outside the infection site, including colonisation of the top of the dermal surface. (Fig. 2D (3b).

### Transcriptomic analysis reveals a pleiotropic regulatory role for SadB

To determine the transcriptional impact of deleting or constitutively expressing *sadB*, RNAseq was performed on total RNA extracted from the WT, Δ*sadB* and *sadB*^+^ (PAO1 (pSadB); Table S1) strains cultured planktonically with shaking at 37°C in LB at both mid-exponential and stationary phase. Following statistical validation of the dataset obtained, only genes with a fold change ≥ ±2 and an adjusted P value of ≤0.05 were considered.

Analysis of differentially regulated genes revealed that a significant proportion of the transcriptome was affected (Fig. 3). Comparison of Δ*sadB* with the WT grown to exponential phase revealed differential regulation of ∼24% of the genome (659 up-regulated genes; 762 down-regulated genes; Fig. 3). In stationary phase ∼8% were differentially regulated (242 up-regulated genes; 210 down-regulated genes). Constitutive expression of *sadB* in the *sadB*^+^ strain compared with the parent resulted in differential expression of ∼2% of the genome in exponential phase (92 up-regulated genes; 37 down-regulated genes). However, in stationary phase, ∼17% of the genome was differentially regulated (558 up-regulated genes; 408 down-regulated genes) (Fig. 3). Of the 648 genes up-regulated in Δ*sadB* in log phase, only 20 remained up-regulated in stationary phase. Of the 762 genes down-regulated in Δ*sadB* in log phase, only 210 genes remained down-regulated in stationary phase (Fig. 3) suggesting a major regulatory role for SadB in planktonic culture during the exponential phase of growth. The key differentially regulated genes are summarized in Table S2.

**FIG 3.**
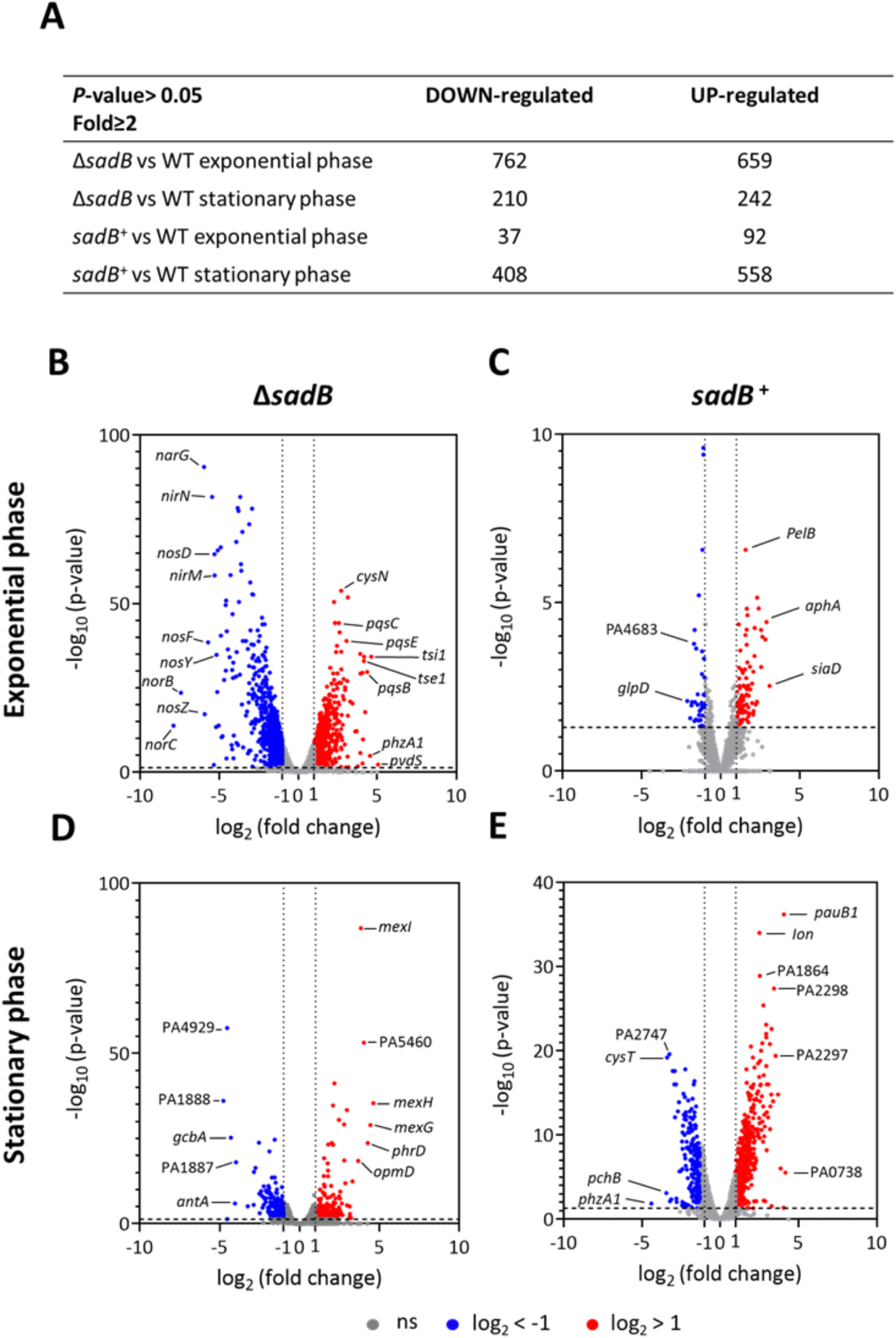
Comparison of the transcriptomes of *P. aeruginosa ΔsadB* and *sadB^+^* with WT in exponential and stationary phase. A) Total number of genes up- and down-regulated. (B-E) Volcano plots of the differentially expressed *ΔsadB* (B, D) and *sadB^+^* (C, E) genes compared with the WT during exponential (B, C) and stationary phases (D, E) of growth. Up-regulated genes are indicated in red (log_2_ fold change>1) and downregulated genes in blue (log_2_ fold change<-1) while grey indicates no significant changes. Statistical cut-off: *p*-value<0.05.

As anticipated, many of the genes at least 2-fold differentially down-regulated in the Δ*sadB* mutant are involved in biofilm development (Table S2). These include genes coding for c-di-GMP generation (e.g the DGCs, SadC and GbcA), c-di-GMP receptors (e.g. *pilZ*), flagella (e.g. *fliC, motB* and *flgM*), pili (e.g. *pilA* and *cupA1*) and expolysaccharides (e.g. *pslA*). Overall, the RNAseq data revealed that diverse genes involved in virulence, protein secretion (type II, II and VI systems), iron and haem acquisition and storage, transcriptional and post-transcriptional gene regulation, quorum sensing (QS), primary and secondary metabolism and respiration were differentially regulated via SadB during log phase planktonic growth when compared with WT (Table S2). The most down-regulated genes in log phase Δ*sadB* were those involved in denitrification (*nar, nir, nor* and *nos*) (Fig. 3B and Table S2).

When *P. aeruginosa* is grown in iron-rich conditions, the *pch* and *pvd* genes required for pyochelin and pyoverdine siderophore synthesis and transport, genes coding for outer membrane receptors for exogenous catechol and hydroxamate siderophore uptake as well as the *has* and *hxu* genes required for heme acquisition are repressed via Fur to prevent iron toxicity (21; 22). Conversely, Fur positively regulates the genes coding for the bacterioferritin (*bfrB*) and ferritin (*ftnA*) iron storage proteins via negative regulation of the PrrF sRNAs. Consistent with these data, *fur, brfB* and *ftnA* were downregulated in the Δ*sadB* mutant in log phase while many of the *pch*, *pvd*, *has* and *hxu* genes as well as *prrF2* were up-regulated suggesting a role for SadB in iron metabolism (Table S2). In agreement with the RNAseq data, deletion of *sadB* resulted in a statistically significant increase in total siderophore production and specifically with respect to pyoverdine (Fig. 4).

**FIG 4.**
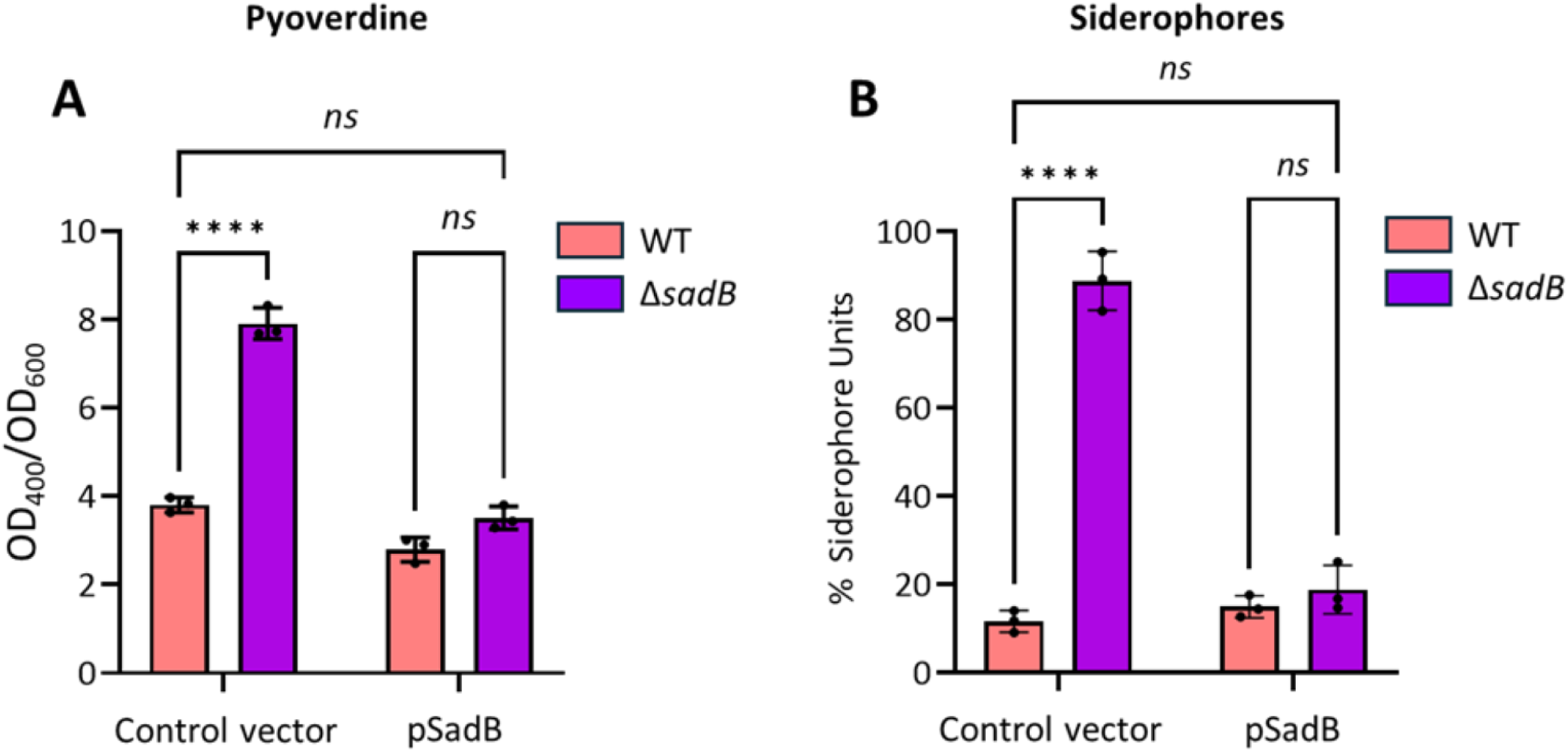
SadB negatively regulates siderophore production. (A) Pyoverdine production measured by determination of the OD_400_ in cell-free spent culture supernatants of the WT and Δ*sadB* mutant strains harbouring the control vector pME6032Δ*lacIQ* or pSadB grown in CAA medium at 37 °C for 24 h. (B) Total siderophore production measured using the CAS method from the supernatants of the four strains tested grown in CAA medium at 37 °C for 24 h. Values given are averages from at least three different cultures ± standard deviation. Statistical differences between group means were determined by two-way ANOVA analysis using Tukey’s multiple comparisons test (\*\*\*\**p* < 0.0001).

The increased expression of the *pqs* genes (Fig. 3B and Table S2) observed in Δ*sadB* is in agreement with the PrrF-dependent enhancement of 2-heptyl-3-hydroxy-4-quinolone (PQS) production (23). This occurs through repression of the anthranilate degradation genes (*antABC*) such that anthranilate can be re-directed towards synthesis of 2-alkyl-4-quinolones (AQs) including PQS. Down-regulation of the *antABC* genes in Δ*sadB* (Fig. 3D and Table S2) is therefore consistent with the PrrF-dependent down-regulation of anthranilate degradation (23).

When *sadB* was constitutively over-expressed in the WT (here termed *sadB*^+^), the most highly up-regulated genes in log phase cells included the biofilm-associated genes *cdrA* and *cdrB*, the *psl* and *pel* exopolysaccharide biosynthesis genes and the *siaABCD* signalling network (Table S2). Examples of genes up-regulated in Δ*sadB* in log phase but down-regulated in the *sadB*^+^ in stationary phase included those involved in iron acquisition (e.g. *pch*, *pvd* and *hxu*) and genes regulated via QS including *phzA1*, *lasB* and *hcnABC* (Table S2).

When collectively considered, the RNAseq data obtained suggests that SadB has a pleiotropic effect on transcription. Deletion of *sadB* had the greatest impact on log phase planktonic cells whereas when constitutively up-regulated in *sadB*^+^, many more genes were differentially regulated in the stationary phase compared with log phase (Fig. 3 and Table S2).

### SadB and c-di-GMP signalling

RNAseq analysis revealed that, in log phase planktonic cells, deletion of *sadB* resulted in the ≥2-fold down-regulation of 9/23 genes coding for c-di-GMP DGCs and 5/12 genes that act as c-di-GMP receptors but with less impact on genes coding for proteins containing EAL or HD-GYP (3/13) domains that act as PDEs (Table S2). The most down-regulated DGC gene in log phase Δ*sadB* was *gcbA* (14 fold) which was further reduced in the stationary phase (19.7 fold). In PAO1, GcbA is involved in the initial stages of surface attachment and influences flagellum-driven motility by suppressing flagellar reversal rates in a manner independent of viscosity, surface hardness, and exopolysaccharide production (24). The *gcbA* RNAseq data for *gcbA* was validated using RT-qPCR which confirmed the down-regulation in log phase when Δ*sadB* was compared with the parent strain but highly up-regulated in *sadB*^+^ in stationary phase (Fig. 5A and B).

**FIG 5.**
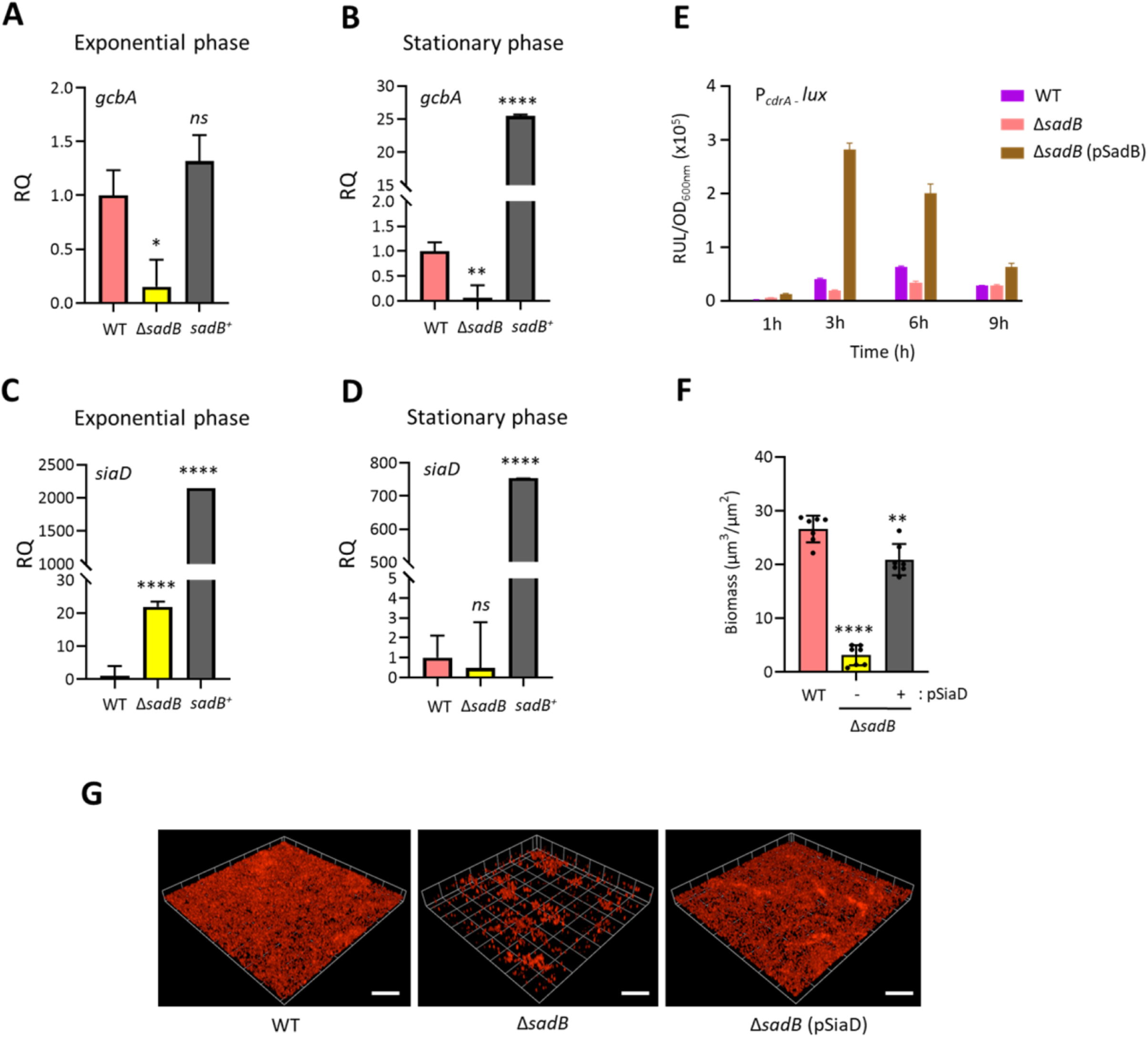
SadB modulates c-di-GMP signalling. (A-D) shows that the DGCs, *gcbA* and *siaD* are differentially expressed in *ΔsadB* and *sadB^+^* compared with WT at both exponential (A and C) and stationary (B and D) growth phases in LB at 37 °C and quantified by RT-qPCR. (E) Indirect quantification of c-di-GMP made using a chromosomally integrated *cdrA’-lux* promoter fusion in WT carrying pME6032Δ*lacIQ*, *ΔsadB* carrying pME6032Δ*lacIQ* and Δ*sadB* carrying pSadB grown in LB at 37 °C in LB. Values correspond to the averages from at least three different cultures ± standard deviation plotted as relative light units normalized to culture density (RLU/OD_600_). (F) Quantification of biofilm biomass from confocal images of WT (pME6032Δ*lacIQ*) and Δ*sadB* harbouring the control vector pME6032Δ*lacIQ* and Δ*sadB* (pSiaD). Values given are averages from at least three different cultures ± standard deviation. (G) Confocal biofilm images of *P. aeruginosa* WT (pME6032Δ*lacIQ*) and Δ*sadB* with or without pSiaD and stained with FM4-64 dye after 48 h incubation in RPMI-1640 medium. Scale bar: 100 µm. Statistical differences between group means were determined by one-way ANOVA analysis using Tukey’s multiple comparisons test to the WT (\**p* < 0.05, \*\**p* < 0.01, \*\*\*\**p* < 0.0001).

The SiaABCD signalling network controls cell aggregation and biofilm formation in *P. aeruginosa* and is important for producing c-di-GMP via SiaD and cell-associated Psl in planktonic cells (26). While no differential regulation of the *sia* operon was noted in the RNAseq for Δ*sadB*, the *sia* genes including *siaD* were all highly upregulated in log phase *sadB*^+^ (Table S2). Similar results were obtained using RT-qPCR for both log (Fig. 5C) and stationary phase (Fig. 5D) *siaD*.

These data highlighted a role for *sadB* in regulating a subset of genes involved in c-di-GMP production and sensing (Table S2). To investigate the impact of *sadB* deletion on c-di-GMP levels as a function of growth, we introduced a *cdrA’::lux* reporter gene fusion that provides an indirect measurement of c-di-GMP levels (27) into the parent, Δ*sadB* and the genetically complemented Δ*sadB* (Δ*sadB* pSadB) strains. Fig. 5E shows that constitutive expression of *sadB* in planktonic Δ*sadB* resulted in a marked rise in light output within the first 3 h consistent with a substantial increase in c-di-GMP production. Lower levels of c-di-GMP were observed in Δ*sadB* compared with PAO1 WT over the first 6 h of growth (Fig. 5E) consistent with (16). Introduction of plasmid borne *siaD* into Δ*sadB* restored biofilm formation Fig. 5F and G). These data establish a clear link between SadB and c-di-GMP signalling in *P. aeruginosa*.

### SadB and the biofilm extracellular matrix

Three exopolysaccharides, Psl, Pel and alginate are associated with the biofilm matrix in *P. aeruginosa* PAO1. Although alginate is primarily associated with the conversion to mucoidy, there is little alginate present in non-mucoid *P. aeruginosa* PAO1 biofilms (28). RNAseq revealed that most of the *psl* operon genes were down-regulated in log phase in Δ*sadB* but up-regulated in the log phase of the *sadB*^+^ strain consistent with the biofilm phenotypes of the respective strains (Table S2 and Fig. 1B). These data suggest that SadB positively influences *psl* expression and were validated using a CTX::*pslA’-luxCDABE* fusion and RT-qPCR of *pslB* (Fig.6A and B). In microtitre plate assays, for both WT and *sadB*^+^ strains but not Δ*sadB*, *pslA* expression peaked in log phase and was highest in *sadB*^+^ (Fig. 6A). The expression of Δ*pslB* was lowest in Δ*sadB* as quantified using RT-qPCR (Fig. 6B). Other biofilm associated genes that were differentially regulated included the *pel* and *cdrA* operons (up-regulated in *sadB*^+^ log phase cells), the chaperone usher pilus gene *cupA1* (down-regulated in Δ*sadB* but up-regulated in *sadB*^+^) and *endA*, an exonuclease involved in eDNA degradation and biofilm dispersal (downregulated in log phase Δ*sadB*) (Table S2).

**FIG 6.**
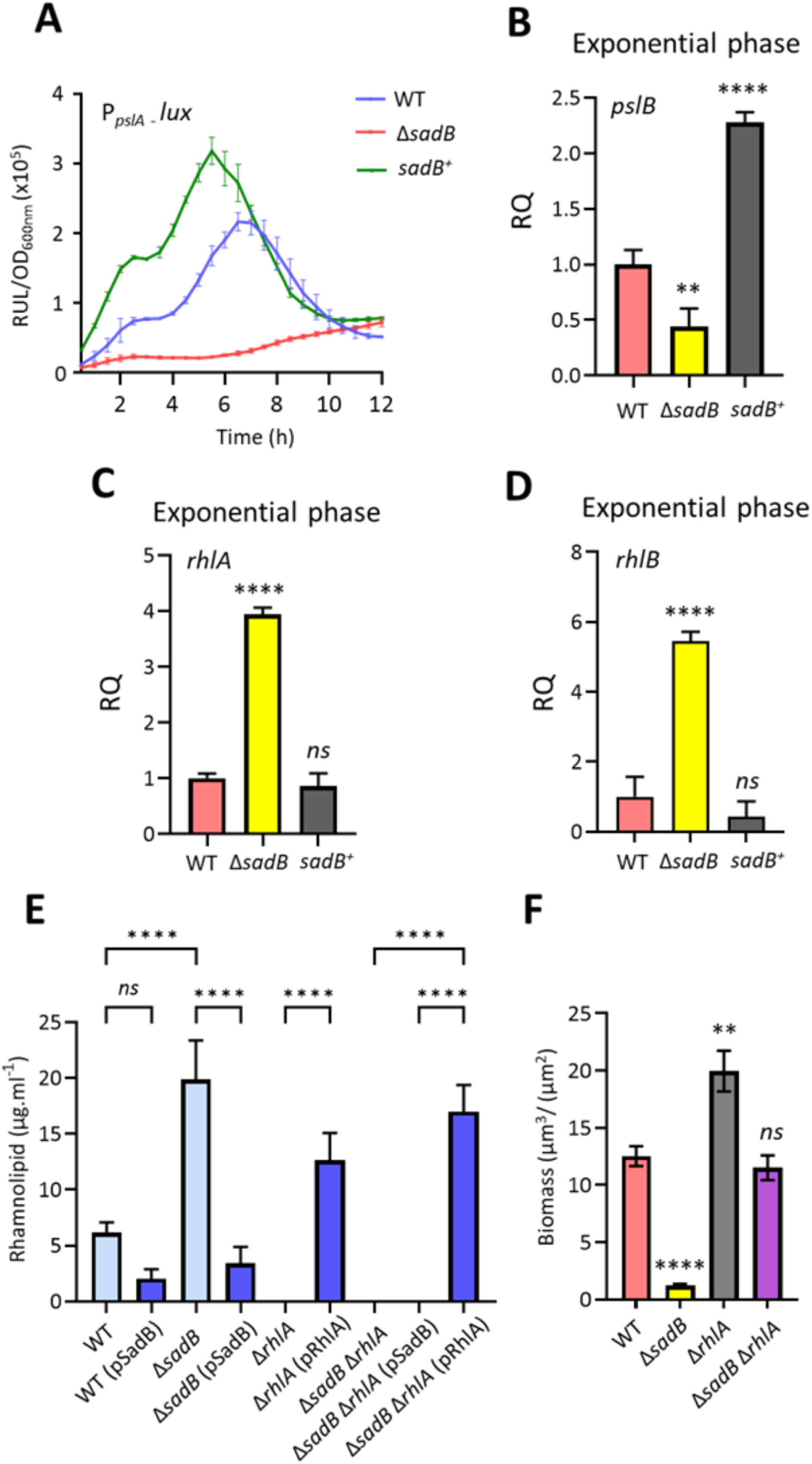
SadB reciprocally regulates genes coding for the exopolysaccharide Psl and the rhamnolipids. (A) Transcriptional activity of a *pslA’*::*lux* promoter fusion in WT and Δ*sadB* mutant with or without pSadB in LB at 37 °C. Values correspond to the averages from three different cultures ± standard deviation plotted as relative light units normalized to cell culture density (RLU/OD_600_). Differentially expressed *pslB* (B), *rhlA* (C) and *rhlB* (D) genes of Δ*sadB* mutant and *sadB^+^* overexpressing strains compared to WT during exponential phase of growth at 37 °C in LB quantified using RT-qPCR. (E) Rhamnolipid production qunatified using the Orcinol method from the cell-free supernatants prepared from WT, Δ*sadB*, Δ*rhlA*, Δ*sadB*Δ*rhlA* and the genetically complemented strains after growth in LB at 37 °C for 24 h. (F) Quantification of biomass from confocal images of WT, Δ*sadB*, Δ*rhlA*, and Δ*sadB*Δ*rhlA* mutants. Values represent the means of at least three biological replicates. Statistical differences between group means were determined by one-way ANOVA analysis using Tukey’s multiple comparisons test to the WT (\*\**p* < 0.01, \*\*\*\**p* < 0.0001).

*P. aeruginosa* produces rhamnolipid surfactants which make multiple contributions to biofilm maturation and architecture ranging from microcolony and channel formation to detachment and dispersal (52, 29). Rhamnolipids are also essential for surface swarming migration. Although deletion of *sadB* in strains PA14 (9) and PAO1 (this manuscript) results in a biofilm defective, hyper-swarming phenotype, the impact of *sadB* on the expression of genes coding for rhamnolipid biosynthesis (*rhlAB*) and hence rhamnolipid production has not, to our knowledge been previously evaluated.

The expression of *rhlAB* was higher in the Δ*sadB* mutant compared with the WT (*rhlA* 6.8-fold, *rhlB* 1.7-fold) in exponential phase (Table S2). RT-qPCR data validated the increased expression of *rhlA* and *rhlB* in the Δ*sadB* mutant (Fig. 6C and D). To quantify the impact of deleting *sadB* on rhamnolipid production, LC-MS/MS was used. The Δ*sadB* mutant produced ∼3.4 and ∼9 fold more than the WT and *sadB^+^* respectively (Fig. 6E). Genetic complementation of Δ*sadB* reduced rhamnolipid production by ∼6-fold. In *sadB^+^,* rhamnolipid production was ∼2.8 -fold lower than in the WT (Fig. 6E).

Since rhamnolipids are biosurfactants that contribute to both biofilm maturation and swarming motility, their overproduction in *ΔsadB* could explain both the biofilm defective and hyper-swarming phenotypes of this mutant. To explore this possibility, we deleted the *rhlA* gene in Δ*sadB.* Fig. 6F shows that biofilm formation could be restored to WT levels in the Δ*sadB*Δ*rhlA* double mutant. Given the reciprocal regulatory links between exopolysaccharides and rhamnolipids via c-di-GMP signalling, we investigated c-di-GMP levels in a Δ*rhlA* mutant compared with Δ*sadB* and a Δ*sadB*Δ*rhlA* double mutant. Fig. S5A shows that c-di-GMP levels are greatly elevated in Δ*rhlA* but reduced back to WT levels in Δ*sadB*Δ*rhlA* consistent with the positive contribution of SadB to the c-di-GMP signalling network. In further support of the role of SadB in biofilm formation, *pslA* expression was highly up-regulated in Δ*rhlA* but reduced back to WT levels in a Δ*sadB*Δ*rhlA* double mutant (Fig. S5B).

### SadB and quorum sensing

*P. aeruginosa* employs a quorum sensing cascade integrating two *N*-acylhomoserine lactone systems (*las* and *rhl*) with an alkylquinolone (*pqs*) QS system (1). Since rhamnolipid biosynthesis is controlled via both the *rhl* and *pqs* QS systems (30), we explored the impact of SadB on QS in the RNAseq dataset. When compared with the WT and in common with the QS target genes, the *N*-butanoyl homoserine lactone (C4-HSL) synthase gene, *rhlI* and the AQ biosynthesis genes (*pqsABCDE phnAB* and *pqsH*) were all highly up-regulated in log phase *ΔsadB* (Table S2) consistent with a loss of population density dependency. Although the *lasRI* genes were not differentially regulated, the gene coding for RsaL, the homeostatic negative regulator of *las*-dependent QS (31) was down-regulated in log phase Δ*sadB*.

To validate the RNAseq data for QS systems, we quantified the production of QS signal molecules in stationary phase culture supernatants using LC-MS/MS. Fig.7 shows that, compared with WT and *sadB*^+^, deletion of *sadB* resulted in statistically significant elevated levels of the AQs, PQS and 2-heptyl-4-hydroxyquinoline *N*-oxide (HQNO) and the RhlI-dependent, short chain *N*-acylhomoserine lactone, *N*-butanoylhomoserine lactone (C4-HSL). However, the difference between WT and Δ*sadB* for the LasI autoinducer*, N*-(3-oxo-dodecanoyl)homoserine lactone (3-oxo-C12-HSL) was not significant (Fig. 7C).

**FIG 7.**
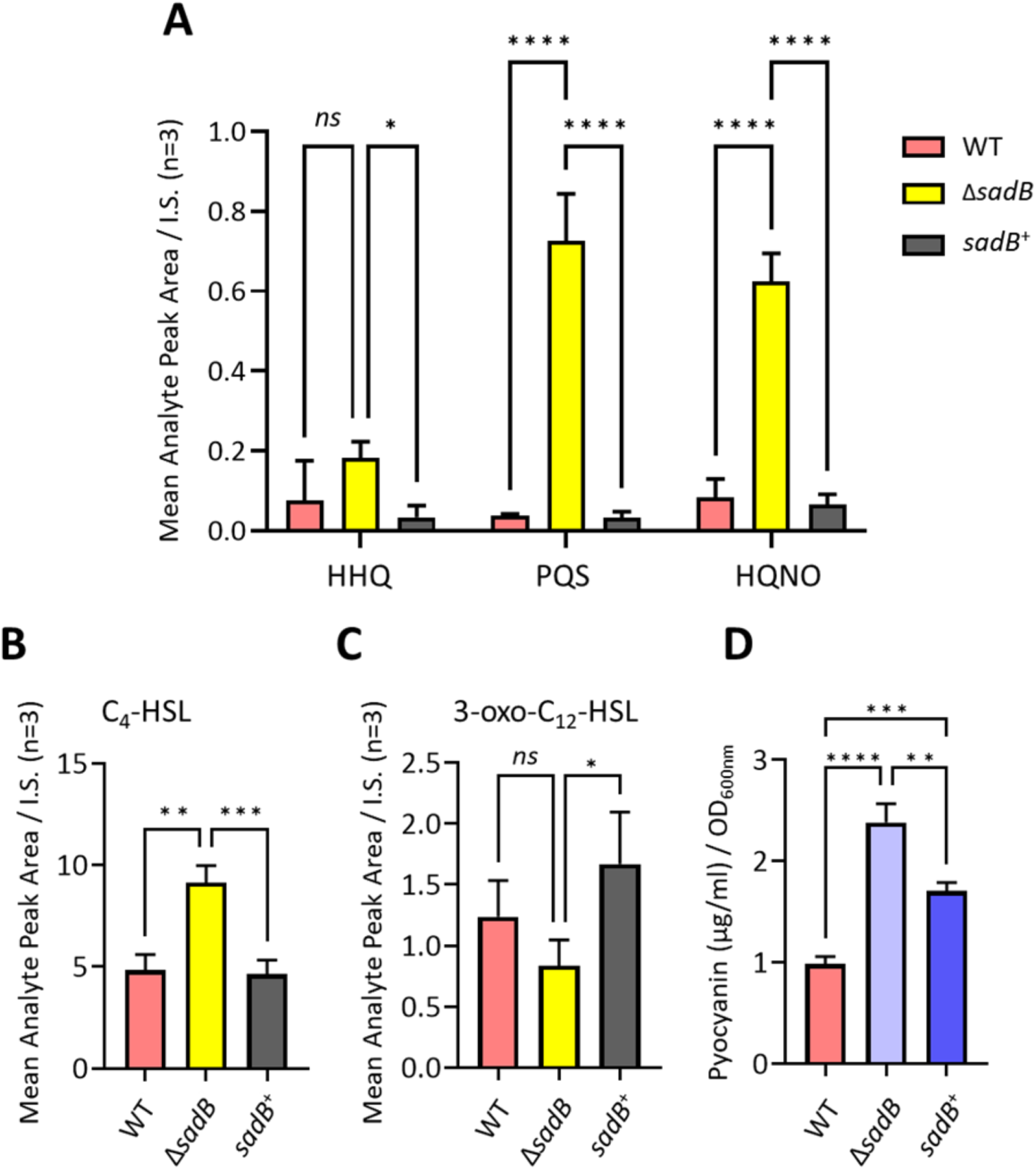
SadB positively regulates AQ and AHL signal molecule and pyocyanin production. (A) HHQ, PQS and HQNO, (B) C_4_-HSL (C) 3-oxo-C_12_-HSL and (D) pyocyanin. AHLs and AQs were quantified using LC-MS/MS while pyocyanin was quantified spectrophotometrically. Values represent the means of at least three biological replicates. Statistical significance was determined by one-way ANOVA analysis using Tukey’s multiple comparisons test. (\**p* < 0.05, \*\**p* < 0.01, \*\*\**p* < 0.005, \*\*\*\**p* < 0.0001).

AQ biosynthesis requires anthranilate which can be supplied via the anthranilate synthases TrpEG and PhnAB or by the degradation of tryptophan via the kynurenine (*kyn*) pathway (32). Table S2 shows that *phnAB* and *kynA* but not *trpEG* expression was higher in log phase *ΔsadB* in log phase compared with the WT (*phnA* 17.9-fold, *phnB* 7.1-fold; *kynA* 2-fold). These data are consistent with the earlier requirement for anthanilate for AQ production in log phase *ΔsadB*. Conversely, anthranilate can be re-directed to the TCA cycle for energy through degradation via the *ant* and *cat* pathways (33). The expression of the *antABC* genes was significantly down-regulated in stationary phase *ΔsadB* (*antA* -16.5-fold, *antB* -9.5-fold, *antC* -5.6-fold;) but highly up-regulated in stationary phase *sadB*^+^ as was the *cat* operon which is responsible for the degradation of the catechol product arising from AntABC-mediated enzymatic degradation of anthranilate (Table S2). These data are consistent with the expression of *antR*, the positive transcriptional regulator for both the AntABC and CatABC degradation pathways, which was up-regulated in the RNAseq data for the *sadB*^+^ strain (Table S2) and confirmed by RT-qPCR (Fig. S3).

Given the log phase induction of the *rhl* and *pqs* QS systems in *ΔsadB,* we examined the RNAseq data for genes known to be targets of *pqs-* and *rhl-* dependent QS. Table S2 shows that in addition to *rhlAB*, those required for the exoproteases (*lasB*, *lasA*, *aprA and pepB* (PA2939), pyocyanin (*phzA1* and *phzB1*), hydrogen cyanide (*hcnABC*), the MexGHI efflux pump and the iron chelators, pyochelin (*pch*) and pyoverdine (*pvd*) were all up-regulated in log phase Δ*sadB*

To validate the relevant RNAseq data, we examined the stationary phase extracellular protein profile of Δ*sadB* by SDS-PAGE (Fig. S4). The levels of two proteins identified by LC M/S as elastase (LasB) and the leucine amino peptidase (PaAP) appeared to be present in greater abundance in Δ*sadB* compared with WT consistent with the increased expression of the corresponding *lasB* (2.8 fold upregulated) and *pepB* (PA2939; 2.7 fold upregulated)) genes both of which are QS-regulated (34). The *rhl* and *pqs* QS-regulated blue-green phenazine pyocyanin is produced in *P. aeruginosa* via either of the two almost identical *phz1* and *phz2* operons together with *phzM* and *phzS* (35). RNAseq analysis showed increased expression of *phz genes* in *ΔsadB* compared with the WT (*phzA1*, 33.2-fold, *phzB1* 8.7-fold) in exponential phase (Table S2) whereas *phzA1* was down-regulated in the *sadB^+^* in the stationary phase compared with Δ*sadB* (*phzA1*, 21-fold). Quantification of pyocyanin levels after growth in flasks with shaking confirmed the RNAseq data in that significantly more pyocyanin was produced by Δ*sadB* compared with both the WT and *sadB^+^* (Fig. 7D).

### *sadB* and denitrification

In microaerophilic or anaerobic environments, *P. aeruginosa* is capable of dissimilatory nitrate reduction for energy production that depends on four enzymatic complexes to reduce nitrate to nitrite (NarGHI), nitrite to nitric oxide (NirS), nitric oxide to nitrous oxide (NorCB), and, finally, nitrous oxide to dinitrogen (NosZ) (36, 37) In the log phase Δ*sadB* RNAseq dataset, *nar*, *nir*, *nor* and *nos* genes were the most down-regulated (from -237 to -12.3 fold; Table S2). In addition, nitrate reduction and in particular the NarGHI enzyme complex requires a molybdate co-factor generated via the *moaA1B1 moaEDC* gene products (38) which were also negatively regulated in log phase *ΔsadB* (Table S2). In the absence of nitrate, *P. aeruginosa* is able to utilize arginine as an energy source via the *arcDABC* operon that codes for the arginine deiminase pathway, which is inducible under oxygen limitation (39). In common with dissimilatory nitrate reduction, the *arcDABC* operon was substantially down-regulated in log phase Δ*sadB* (Table S2).

To evaluate the consequences arising from the reduced expression of genes required for growth in reduced oxygen, we grew the *P. aeruginosa* WT, Δ*sadB* and *sadB*^+^ strains either statically or with shaking or in microtitre well plates. Under aerobic shaking conditions, all strains grew to similar cell densities (WT, OD_600_ 2.9 ± 0.1; *ΔsadB* OD_600_ 2.9 ± 0.03 and *sadB^+^* OD_600_ 3.0 ± 0.1) (Fig. 8A). In static conditions after 10 h growth, there was a clear reduction in the *ΔsadB* growth (OD_600_ 0.47 ± 0.01) compared with the WT (OD_600_ 0.91 ± 0.04) or *sadB^+^* (OD_600_ 1.4 ± 0.1) (Fig. 8B).

**FIG 8.**
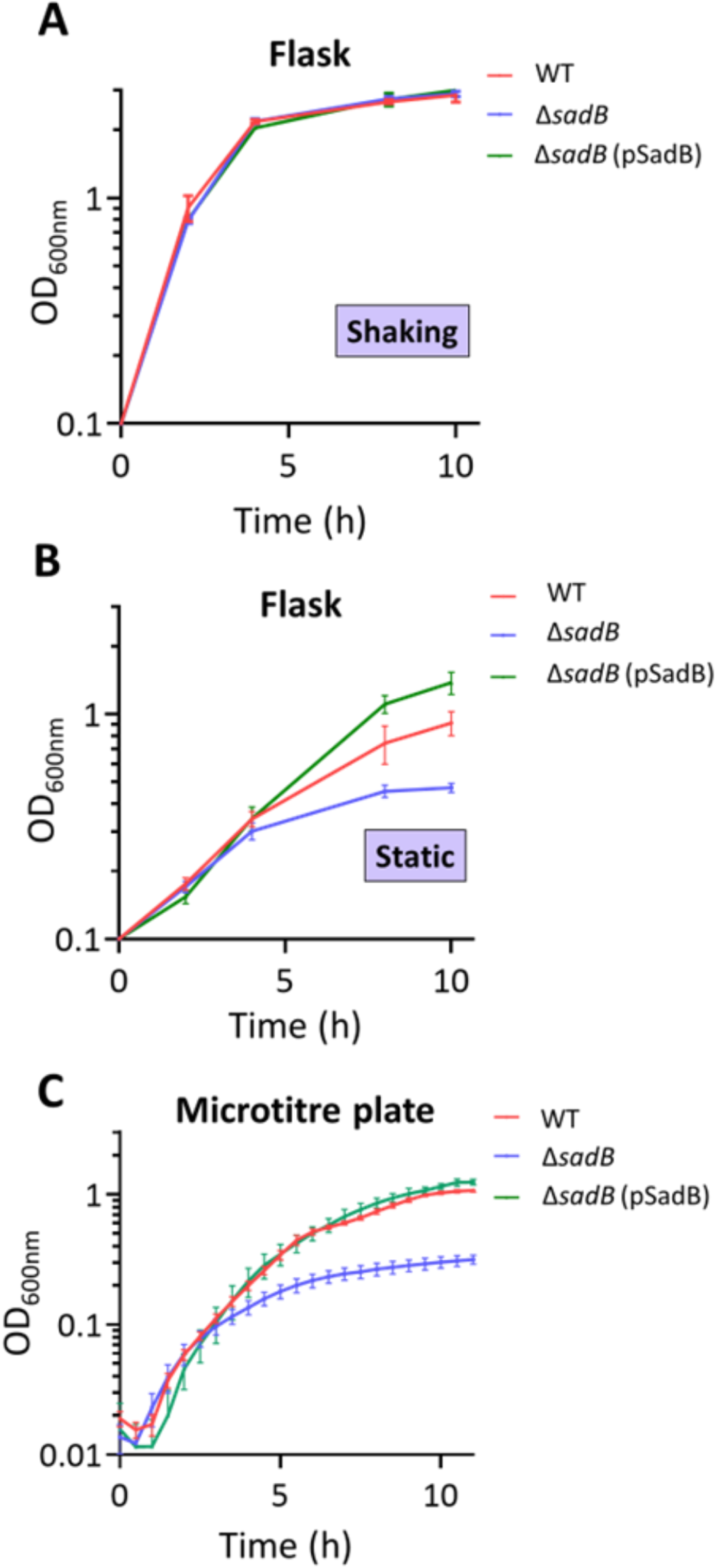
Oxygenation influences the growth of Δ*sadB*. Growth of WT and Δ*sadB* harbouring the control vector pME6032Δ*lacIQ*, and Δ*sadB* pSadB in 50 ml conical tubes for 12 h in LB 37 °C under vigorous shaking (200 rpm) (A) or static (B) conditions or (C) in a 96-well microtitre plate. Values given are averages from at least three different cultures ± standard deviation.

In microtitre plates, the WT and complemented Δ*sadB* grown statically reached stationary phase and their maximum OD_600_ at ∼10 h (Fig. 8C). A growth defect was clearly apparent for Δ*sadB* in which growth began to slow between 3 and 4 h. Since down regulation of the denitrification genes in log phase *ΔsadB* may be responsible for the microaerophilic growth defect, we supplemented LB with a range of nitrate concentrations. Fig. S6 shows a small increase in Δ*sadB* growth under microaerophilic conditions when provided with 3.2 mM KNO_3_.

## DISCUSSION

The major aim of this work was to obtain new insights into the contribution of *sadB* to *P. aeruginosa* virulence and biofilm formation and to elucidate its downstream targets. In an acute mouse skin and soft tissue infection model, D*sadB* was unable to establish infection and was rapidly cleared from the infection site. This contrasted with both the WT parent and genetically complemented D*sadB* both of which established within the host tissues forming architecturally distinct infection sites. The higher luminescence output (∼1.5 log fold) from the genetically complemented D*sadB* CTX::*tac’-sadB-luxCDABE* strain reflected an increased population of metabolically active bacteria on 3 and 4 days after inoculation. In addition, the greater size of the infection site, lack of a fibrotic pocket and marked infiltration of leukocytes into both infection site and into the adipose and subdermal tissue layers further distinguished between WT and D*sadB* CTX::*tac’-sadB-luxCDABE* strain indicating that the latter was more virulent. A PA14 *sadB* Tn insertion mutant has previously been reported to be attenuated in a rat model of chronic lung infection (40). These data are consistent with a broader role for SadB in the pathogenesis of *P. aeruginosa* infections than previously appreciated.

To evaluate the global impact of *sadB* deletion and to gain further insights into the reduction in virulence observed, we undertook RNAseq analysis of planktonic cells, firstly by comparing WT with D*sadB* grown to either exponential or stationary growth phase. The data obtained surprisingly revealed that deletion of *sadB* resulted in the differential regulation of over 1420 genes by ≥ ±2 fold in log phase D*sadB* compared with WT. Far fewer genes were differentially regulated (∼450 ≥ ±2 fold) by the time the cells reached stationary phase. Of particular interest, was the log phase induction of genes associated with the *P. aeruginosa rhl* and *pqs* QS systems in D*sadB* indicating that SadB is a negative regulator of QS. Within the *pqs* system, PQS acts as an extracellular QS signal molecule and iron chelator that autoinduces AQ biosynthesis as well as the expression of genes involved in the iron-starvation response and virulence factor production via PqsR-dependent and PqsR-independent pathways (41). Given the impact of *sadB* deletion of iron metabolism, it is interesting to note that impairment of iron release from bacterioferritin (BfrB), which is downregulated in log phase D*sadB*, has been reported to result in reduced biofilm formation but increased rhamnolipid production and swarming (42).

The dispensable *pqs* system thioesterase, PqsE, is involved in the regulation of diverse genes coding for key virulence determinants and biofilm development via a direct interaction with RhlR, the C4-HSL-dependent regulator (43). Although *rhlR* was not differentially regulated in log phase Δ*sadB,* the C4-HSL synthase gene *rhlI* and the *rhlA* and *rhlB* genes required for rhamnolipid biosynthesis were all up-regulated consistent with the higher levels of PQS, C4-HSL and rhamnolipids quantified in stationary phase Δ*sadB.* The log phase induction of the *pqs* QS system is also in accordance with the increased expression of genes involved in high affinity iron transport systems, the down-regulation of anthranilate degradation and the substantially reduced expression of the *nir*, *nar* and *nos* denitrification genes. The pivotal role of QS in the adaptation of *P. aeruginosa* to anaerobic growth is well established in the literature (44, 45, 46).

Since QS systems are, by definition, cell population density dependent, their log phase induction in Δ*sadB* may alert host defences to the presence of low numbers of infecting bacteria and their virulence factors and so promote clearance before the pathogen becomes established. However, the log phase induction of QS in Δ*sadB* is unlikely to provide a full explanation given that QS, is just one of several different environmental parameters (e.g. temperature, pH, osmolarity, oxidative stress, nutrient deprivation) which bacterial cells must integrate to determine their optimal survival strategy within a given environment (47).

In microaerophilic but not aerobic conditions, Δ*sadB* exhibited a growth defect. Supplementation with nitrate in contrast to the genetic complementation of *sadB* only partially increased growth under microaerophilic conditions consistent with the down-regulation of both the denitrification and molybdenum co-factor genes in Δ*sadB.* This could also conceivably compromise *P. aeruginosa* growth *in vivo* where leucocyte-dependent utilization of oxygen to generate antibacterial free radicals in the tissues can cause localized hypoxia (25). In support of these observations, *sadB* has been reported in PAO1 to be expressed more highly under both low oxygen (48) and anaerobic conditions (49). These findings suggest that *sadB* is likely important for energy generation in oxygen limiting conditions and for the normal operation of the denitrification pathway. For both PA14 and PAO1, *sadB* mutants are hyper-swarmers that fail to form biofilms. The PA14 *sadB* mutant biofilm defect has been associated with an inability to switch from reversible attachment (where the bacterial cells are in a relatively unstable surface contact via their poles) to irreversible attachment where the cells are surface aligned via their long axes (10). Our finding that the rhamnolipid biosynthetic genes are up-regulated in log phase planktonic Δ*sadB* leading to overproduction of rhamnolipids suggested a further explanation of the Δ*sadB* biofilm defect. Rhamnolipids are multifunctional biosurfactants that have surface lubricating, anti-adhesive properties and contribute to biofilm maturation and dispersal (50, 51). Premature overproduction of surface lubricating rhamnolipids by Δ*sadB* is therefore likely to account for the inability of *P. aeruginosa* to switch from reversible to irreversible surface attachment. This suggestion is supported by our finding that deletion of *rhlA* in Δ*sadB* results in the restoration of biofilm formation with the double mutant forming a biofilm of similar biomass to the WT.

The RNAseq data also revealed that deletion of *sadB* had a major impact on the transcription of many genes involved in c-di-GMP signalling. In PAO1 when c-di-GMP levels were maintained at low levels by constitutive expression of the PDE, YhJH, both *rhl* and *pqs* QS systems were expressed at higher levels (53) leading to increased production of *rhl* and *pqs*-regulated virulence factors including pyocyanin and rhamnolipids. This is consistent with the Δs*adB* data. However, (53) proposed that the induction of QS-regulated virulence factors in cells with low intracellular c-di-GMP levels and especially rhamnolipids would compromise host defences and enhance virulence. Contrary to this, we found that Δ*sadB* was unable to establish infection and was readily clearly from the infection site in a mouse soft tissue infection model indicative of a less virulent phenotype that probably reflects the premature induction of QS and the pleiotropic impact of *sadB* on the transcriptome. Furthermore, the constitutive over-expression of *sadB* resulted in higher levels of cdi-GMP and a more persistent phenotype in our mouse infection model.

Apart from deleting *rhlA*, biofilm formation by Δ*sadB* could also be restored by expressing the DGC gene *siaD* which was highly upregulated in log phase *sadB*^+^. The *sia* system maintains low level c-di-GMP and Psl in planktonic *P. aeruginosa* such that *sia* mutants exhibit a surface attachment-deficient phenotype (26). Overexpression of *siaD* in Δ*sadB* resulted in increased c-di-GMP levels as indicated by the elevated expression of the *cdrA’*::*lux* fusion. Given the role of the *sia* operon in up-regulating the production of Psl, which functions as an early stage adhesive, extracellular biofilm matrix component and signal molecule (54), it is also possible that the restoration of biofilm formation in Δ*sadB* by SiaD is also a consequence of increased Psl production. This is possibly via c-di-GMP-dependent activation of Psl synthesis and also through diversion of biosynthetic substrates from rhamnolipid synthesis. Competition for common sugar precursors catalyzed via AlgC has been proposed as a mechanism for balancing the synthesis of Psl and rhamnolipids so helping to co-ordinate the inverse control of swarming motility and biofilm formation (55, 56). Consistent with sugar precursor competition, we observed that both c-di-GMP levels and *psl* expression are both significantly up-regulated in a Δ*rhlA* mutant but reduced back to WT levels in the Δ*sadB* Δ*rhlA* double mutant highlighting the importance of SadB in this context. Furthermore, the c-di-GMP PDE, NbdA plays a key role diverting of substrates from Psl to rhamnolipids (55, 56). Interestingly, *nbdA* was highly upregulated in stationary phase Δ*sadB* (Table S2).

Although SadB from *P. fluorescens* F113 has been reported to function as a c-di-GMP binding protein (15), structural and c-di-GMP binding studies undertaken by (16) indicate that *P. aeruginosa* SadB, despite its similarity, is highly unlikely to require c-di-GMP or other small molecule ligands for activation. Instead, SadB was demonstrated to bind directly to the C-terminal domain of AmrZ, leading to the rapid proteolytic degradation of this transcription factor primarily via the Lon and to a much lesser extent by the AsrA and PepA proteases (16). Interesting we found that both *lon* and *asrA* genes were significantly upregulated in *sadB*^+^ stationary cells by 5.7 and 2.2 fold respectively.

RNAseq of log phase PAO1 Δ*amrZ* compared with the genetically complemented mutant has revealed 338 differentially regulated genes (at least 2 fold; 89 up; 249 down) (18). Many of these genes are shared with the SadB regulon (1421 2-fold differentially regulated log phase genes; Fig. 3B and 3C) including those involved in iron acquisition, c-di-GMP signalling, quorum sensing, biofilm formation, protein secretion and motility. In the Δ*amrZ* RNAseq dataset (18)*, rhlA* was 2-fold downregulated whereas *rhlA* was highly up-regulated in Δ*sadB* (this manuscript) consistent with the SadB-driven degradation of AmrZ and the respective swarming and biofilm phenotypes. It is therefore clear that while SadB clearly mediates substantial regulatory activity through AmrZ degradation, it may have additional targets given the apparently larger size of its regulon compared with that of AmrZ. Furthermore, the highly attenuated virulence of the *sadB* mutant together with recent developments in the structure and function of SadB suggest that it has considerable potential as a novel protein target for structure-based antibacterial drug discovery.

## MATERIALS AND METHODS

### Bacterial strains, plasmids, oligonucleotides and culture conditions

The bacterial strains, plasmids and oligonucleotides used in this study are listed in Tables S1 and S3. *P. aeruginosa* and *E.coli* strains were routinely cultured in lysogeny broth (LB) at 37°C with shaking at 200 rpm or on LB agar. For some experiments, bacteria were also grown statically in LB with or without a range of potassium nitrate concentrations. Where required the following antibiotics were added: ampicillin (Ap) 100 µg/ml (*E. coli*); carbenicillin (300 μg/ml) (*P. aeruginosa*); tetracycline (Tc) 25 µg/ml (*E. coli*) or 125 µg/ml (*P. aeruginosa*) and nalidixic acid (NA) 30 μg/ml (for *P. aeruginosa* selection in mating experiments with *E. coli*). *P. aeruginosa* swarming assays were carried out on in petri dishes on a solid medium containing Nutrient Broth No. 2 (Oxoid) 8 g/L, D-glucose 0.5% w/v and Bacto agar (0.5% w/v; Difco) as described by (30).

### Construction of *P. aeruginosa* deletion mutants and complementation vectors

*P. aeruginosa* Δ*sadB* and Δ*sadB*Δ*rhlA* deletion mutants were constructed via allelic exchange. Two PCR products amplifying the upstream and the downstream regions of each gene were generated using the primer pairs 1FW/1RW, and 2FW/2RW respectively (Table S3). The resulting PCR products were cloned into the suicide plasmid pME3087 (57) and introduced into *P. aeruginosa* via conjugation with *E. coli* S17-1 λpir. Recombinants were selected on tetracycline followed by enrichment with carbenicillin (58). Deletions were confirmed by DNA sequence analysis and their swarming and biofilm phenotypes confirmed (Fig. 1). The *sadB* and *siaD* expression vectors (pSadB and pSiaD; Table S1) were constructed by PCR-amplification using PAO1 chromosomal DNA as a template and with primer pairs SadB F/R and SiaD F/R respectively. *sadB or siaD* were each cloned into the shuttle vector pME6032 which contains a *lacIQ* mutation rendering the P*tac* promoter constitutive (59). The *sadB* gene was also cloned into the pColdI protein expression vector to generate pMES1 (Table S1) and transformed into *E. coli* BL21. After culturing in Terrific broth, *sadB* expression was induced with IPTG (1 mM) after cold shock and the bacterial cells harvested by centrifugation and lysed. The lysate was subjected to nickel chromatography followed by size exclusion chromatography to purify the His-tagged SadB protein. Polyclonal antibodies against SadB were raised in rabbits,

### Construction of bioluminescent strains for *in vivo* experiments

The miniCTX::*tac’-luxCDABE* promoter fusion described previously (60) was introduced onto the chromosomal CTX site of both WT parent and Δ*sadB.* In addition, a CTX::*tac’-sadB-luxCDABE* expression vector was constructed by PCR-amplification of the *sadB* gene using PAO1 chromosomal DNA as a template and the primer pairs P_tac_sadBlux F/R (Table S3). The resulting PCR product carrying *sadB* gene was ligated into the miniCTX*lux* vector (Table S1) using the appropriate restriction enzyme to generate miniCTX::*tac’-sadB-luxCDABE*. The resulting *lux*-marked *sadB* expression vector was integrated into the chromosome of *P. aeruginosa* by conjugation with *E. coli* S17.1 λpir followed by selection with tetracycline.

### Construction of *pslA’*::*lux* transcriptional reporter fusion

The promoter regions of *pslA* were amplified from the PAO1 genome using the primer pair PpslA’F/R (Table S3). The resulting PCR product was ligated into the mini-CTX*lux* vector using the appropriate restriction enzyme to generate the mini-CTX::*pslA’*-*lux* transcriptional reporter. The transcriptional reporter was integrated into the *P. aeruginosa* chromosome by conjugation with *E. coli* S17.1 λpir followed by selection with gentamicin.

### Mouse infection model

A simple and reproducible mouse model based on that described by (19) was used. All animal experiments were approved following local ethical review at the University of Nottingham and performed under UK Home Office licence PP5768261. Female BALB/c mice, 19–22g were housed in individually vented cages under a 12 h light cycle, with food and water *ad libitum*. Animals were anaesthetised with 2% isofluorane, their flanks shaved and the skin cleaned with Hydrex surgical scrub. *P. aeruginosa* WT and Δ*sadB* strains carrying chromosomally integrated CTX::*tac’-luxCDABE* or CTX::*tac’-sadB-luxCDABE* fusions (1×10^5^ cfu; Table S1) were mixed with Cytodex 1 microcarrier beads (19) 1 μg/50 μL total volume phosphate buffered saline (PBS) pH 7.4 and co-injected subcutaneously. Animals were allowed to recover and monitored throughout the 4 day study. The progress of bacterial infection was tracked by following bioluminescence output using an IVIS™ Spectrum (PerkinElmer) (61). Infected animals were imaged daily for 4 days for the presence of metabolically active bacteria at the infection sites. Mice were humanely euthanized and infection site tissues excised for histological assessment were dissected, fixed and stained with haematoxylin and eosin as described by (61). Bacterial cells were visualized by IHC using polyclonal antibodies raised against *P. aeruginosa* (Invitrogen PA1-73116) and detected with an anti-rabbit Alexa 555 antibody conjugate (Thermofisher). Tissues were counterstained for DNA with POPO-1. Slides mounted with Fluoromount (Sigma-Aldrich). Images were acquired on a Zeiss Axioscan 7 slide scanner, objective 20x.

### RNA extraction, cDNA preparation and RT-qPCR

Bacteria were grown in 5 ml of LB in 50 ml conical tubes at 37°C with shaking at 200 rpm for 19 h. RNA was extracted from mid-log and stationary phase cultures using the RNeasy Mini Kit (QIAGEN) together with RNAprotect Cell Reagent (QIAGEN) to preserve RNA integrity. Residual genomic DNA was digested using a Turbo DNA-free kit (Invitrogen) and PCR to confirm the absence of DNA contamination. RNA quality and quantity were assessed using an Agilent 2100 Bioanalyser and the Agilent RNA The RNA was used as a template for cDNA synthesis using the GoScript™ Reverse Transcriptase kit (Promega, US). qPCR for the relative expression of target transcripts was performed using an ABI 7500 Fast Real-Time PCR System (Applied Biosystems) and Power SYBR™ Green PCR Master Mix according to the manufacturer’s instructions. The oligonucleotides used for qPCR are listed in Table S3. The *rpoS* gene was used as the internal ‘housekeeping’ control to normalize the qPCR data as it exhibited the least variation in expression in the RNAseq dataset. The analysis was performed at least in duplicate on three technical replicates.

### Transcriptome analysis

RNA sequencing and data analysis were carried out by Novogene. Sequencing libraries were generated using NEBNext® Ultra™ RNA. Library Prep Kit for Illumina® (NEB, USA). The RNA-Seq results have been deposited in NCBI’s Gene Expression Omnibus and are accessible through GEO Series accession number GSE302429.

### Biofilm formation and imaging

Biofilms were grown on borosilicate glass coverslips in RPMI-1460 (Lonza) at a temperature of 37 °C for a period of 24 h. Biofilms were stained with FM4-64 dye (5μM) and were visualised using a confocal laser scanning microscope (LSM700, Carl Zeiss) and 555 nm laser. At least 5 replicate Z-stack biofilm images were taken randomly using Zen 2011 imaging software. Biomass was quantified using Image J (NIH, Bethsda MD, USA) and Comstat 2.1. (https://www.comstat.dk/, 62).

### Bioluminescent reporter assays

To evaluate the expression of *cdrA’*::*lux* and *pslA’*::*lux* transcriptional fusions (Table 1) as a function of growth, bacteria were grown in 96 well plates (flat black transparent bottom, GreinerBioOne). OD_600_ and luminescence (Relative Light Units, RLU) measurements were taken every 30 min at 37°C for 24 h using the TECAN Infinite® F200Pro device. Each assay was performed independently at least twice with three technical replicates.

### QS signal molecules, pyocyanin and rhamnolipid quantification

The *N*-acylhomoserine lactones, C4-HSL and 3-OC12-HSL and the AQs (PQS, HHQ and HQNO) were quantified using LC-MS/MS after extraction from culture supernatants with acidified ethyl acetate as described by (63). Pyocyanin was extracted with chloroform prior to LC-MS/MS. Rhamnolipids were quantified by LC-MS/MS after growth in M9 medium containing 2% glycerol, 2 mM MgSO4 and 0.05% glutamic acid at 37°C with shaking at 200 rpm (64).

### Siderophore Assays

Total siderophores present in spent cell-free culture supernatants prepared from *P. aeruginosa* grown in casamino acids (CAA) medium (65) were quantified using the CAS assay (66) with desferrioxamine as a positive control. Pyoverdine was quantified spectrophotometrically as described by (67) by determining the absorbance of supernatants at 405 nm (A_405_) and dividing by OD_600_.

### Extracellular protein profiles

Cell free culture supernatants from *P. aeruginosa* WT, Δ*sadB* and *sadB*^+^ grown to stationary phase were treated with trichloroacetic acid (10% v/v) for 30 min on ice. The precipitated proteins were collected by centrifugation, washed with acetone prior and air dried. Samples were heated at 90°C for 5 min in sample buffer prior to SDS-PAGE. After staining with Coomassie blue, selected bands were excised and sent for MALDI-MS/MS.

### SDS-PAGE and Western blotting

*P. aeruginosa* cytoplasmic fractions were subjected to SDS-PAGE and transferred to nitrocellulose membranes by Western blotting. After blocking with Tris-buffered saline (pH 8) containing 5% w/v skimmed milk powder, the nitrocellulose was probed with a polyclonal rabbit antibody raised against the purified SadB protein followed by HRP-conjugated anti-rabbit IgG and the blots exposed to Hyperfilm™ (GE) after incubation with an enhanced chemiluminescence HRP substrate (Thermo-Fisher).

### Statistical analysis

One-way and two-way ANOVA analysis using Tukey’s multiple comparison tests were applied to determine whether *P. aeruginosa* derivative strains responses differed significantly from that of the parental strain (p < 0.05) when compared with the variations within the replicates (n ≥ 3) using GraphPad Prism 8.0 (GraphPad Software, Inc., San Diego, CA).

## ACKNOWLEDGEMENTS

We thank Nigel Halliday for LC-MS/MS quantification of *P. aeruginosa* QS signals, and rhamnolipids, Philip Bardelang for help with SadB expression and purification and Matthew Fletcher for constructing the *pslA’*::*lux* fusion. This work was supported via Wellcome Trust joint senior investigator awards to PW and MRA (grant nos. 103882 and 103884), a Wellcome Trust doctoral training programme (108876/B/15/Z) and an EPSRC grant (EP/X001156/1).

## DATA AVAILABILITY

The RNA-Seq results have been deposited in NCBI’s Gene Expression Omnibus and are accessible through GEO Series accession number GSE302429.

